# Biofilm structure promotes coexistence of phage-resistant and phage-susceptible bacteria

**DOI:** 10.1101/552265

**Authors:** Matthew Simmons, Matthew C. Bond, Britt Koskella, Knut Drescher, Vanni Bucci, Carey D. Nadell

## Abstract

Encounters among bacteria and their viral predators (bacteriophages) are among the most common ecological interactions on Earth. Further, these encounters are likely to occur predominantly within surface-bound communities that microbes most often occupy in natural environments. These communities, termed biofilms, are spatially constrained such that interactions become limited to near neighbors; diffusion of solutes and particulates is reduced; and there is pronounced heterogeneity in nutrient access and physiological state. It is appreciated from prior, abstracted theory that phage-bacteria interactions are fundamentally different in spatially structured contexts, as opposed to well-mixed liquid culture. Spatially structured communities are predicted to promote the protection of susceptible host cells from phage exposure, and thus weaken selection for phage resistance. The details and generality of this prediction in realistic biofilm environments, however, are not known. Here we explore phage-host interactions using experiments and simulations that are tuned to represent the essential elements of biofilm communities. Our simulations show that in biofilms, the coexistence of susceptible and phage-resistant bacteria is highly robust to a large array of conditions, including background growth rate, cost of phage resistance, mechanism of phage resistance, and phage diffusivity. We characterize the population dynamics underlying this coexistence, and we show that coexistence is recapitulated in an experimental model of biofilm growth measured with confocal microscopy. Our results provide a clear view into the dynamics of phage-resistance in biofilms, with single-cell resolution of the underlying cell-virion interactions, linking the predictions of canonical theory to realistic models and *in vitro* experiments of biofilm growth.

**Importance:** In the natural environment, bacteria most often live in communities bound to one another by secreted adhesives. These communities, or biofilms, play a central role in biogeochemical cycling, microbiome functioning, wastewater treatment, and disease. Wherever there are bacteria, there are also viruses that attack them, called phages. Interactions between bacteria and phages are likely to occur ubiquitously in biofilms. We show here, using simulations and experiments, that biofilms will almost universally allow phage-susceptible bacteria to be protected from phage exposure, if they are growing alongside other cells that are phage-resistant. This result has implications for the fundamental ecology of phage-bacterial interactions, as well as the development of phage-based antimicrobial therapeutics.

## Introduction

Because of the sheer number of bacteria and phages in nature, interactions between them are very common (1–9). The imperative of evading phages on the part of their bacterial hosts – and of accessing hosts on the part of phages – has driven the evolution of sophisticated defensive and offensive strategies by both (10, 11). Phage resistance can evolve very rapidly in well-mixed liquid cultures of bacteria under phage attack (2, 12, 13); for spatially structured environments, on the other hand, recent work has suggested that selection for phage resistance can take on very different forms due to protection of phage-susceptible cells in confined refugia (14–17). The generality of this prediction in realistic biofilm conditions is currently unknown; here we leverage a custom biofilm-specific simulation framework and a microfluidics-based experimental system to address this question.

Biofilms are characteristically heterogeneous, including steep gradients in nutrient availability, waste product accumulation, oxygenation, and pH, among other factors (18, 19). Furthermore, biofilm structure can impede the movement of solutes and particles that ordinarily would pose grave threats in well-mixed liquid conditions. The extracellular matrix of *Pseudomonas aeruginosa*, for instance, can block the diffusion of antibiotics such as tobramycin (20, 21). Biofilm matrix secreted by *Escherichia coli* and *P. aeruginosa* can also alter phage movement (22, 23), and mucoid colony phenotypes, which correlate with higher capsule or matrix secretion, rapidly evolve under lytic phage exposure in *E. coli* and *P. fluorescens* (24, 25).

Beyond their deep importance to microbial natural history, phages’ ability to rapidly destroy susceptible populations makes them attractive as alternative antimicrobials (12, 26, 27). Optimizing phages for this purpose, including an understanding of phage resistance evolution among host bacteria, requires exploration of models and experiments that specifically capture the essential elements of biofilm environments (28, 29). In particular, biofilm growth may have profound impacts on the relative advantages and disadvantages of phage resistance, because the spatial structure within biofilms can potentially protect susceptible cells from phage exposure (15, 17, 22, 23, 30, 31).

Here we use high resolution imaging of our experimental biofilm system to address this problem, exploring the population dynamics of phages, sensitive bacteria, and resistant bacteria. Our observations indicate that sensitive bacteria coexist at high densities alongside resistant bacteria because of the protection afforded by spatial structure: the resistant bacteria block phage access to sensitive cells and can even act as phage sinks. Computational studies developed in parallel to the experiments support this interpretation and indicate that the protection of sensitive cells generalizes robustly to a wide range of bacterial resistance mechanisms, fitness cost of phage resistance, baseline bacterial growth rates, and diffusivity of phages inside and outside of the biofilm microenvironment.

## Results

In biofilm environments, the population dynamics of bacteria and their lytic phages are driven by many processes, including bacterial growth, cell-cell shoving, solute advection/diffusion, phage-host attachment probabilities, phage lag time and burst size, and phage advection/diffusion, among others (9, 15). To study these processes we expanded a simulation framework previously developed by our groups that captures the biological and solute/particle transport processes inherent to biofilm communities (15) (SI Materials and Methods; code available at https://github.com/simbiofilm/simbiofilm (32)). Our framework implements the growth of up to hundreds of thousands of discrete bacteria and phages in explicit space; it is custom-made for this application but falls within the family of biofilm simulation techniques that have been highly successful in capturing the qualitative dynamics of natural systems (33–35). To begin a simulation, cells are inoculated onto a solid surface at the bottom of a 2-D space with lateral periodic boundary conditions. Growth-limiting nutrients diffuse from a bulk liquid layer at the top of the 2-D space towards the biofilm front, where they can be depleted due to consumption by cells (Figure 1A). The biofilm surface erodes in a height-dependent manner, reflecting the increase in shear rate with distance from the surface (36). After a pre-set interval of biofilm growth, phages are introduced to the system in a pulse at one location along the biofilm’s upper surface (varying the timing or location of phage pulses had little impact on the results, see Supplementary Information). In the simulations, phages can associate with cells in the biofilm and initiate infections, or be released into the surrounding liquid, where they diffuse for a full simulation iteration cycle prior to being swept out of the system by advection (Figure 1A). We implemented phage diffusion by algorithmic rules that are described in detail in the SI Materials and Methods.

**Figure 1.**
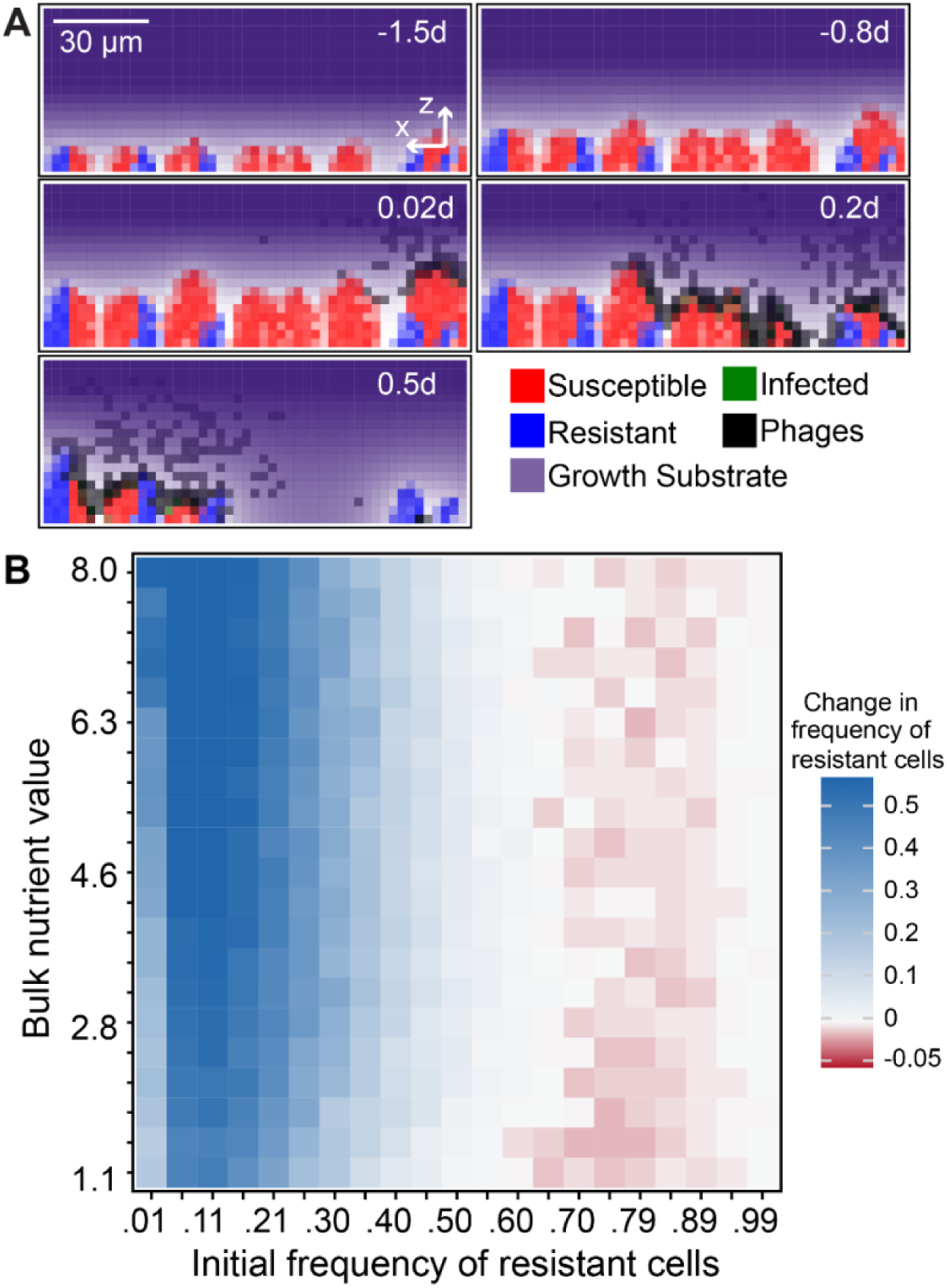
Simulated outcomes of phages exposed to biofilms composed of resistant and susceptible cells. (A) Example time series in which biofilms of phage-resistant and phage-susceptible cells are allowed to reach a critical height before introduction of phages at one location along the biofilm surface (varying the initial biofilm height and phage introduction procedure are explored in Figure S3). Phages can absorb to resistant cells but cannot amplify within them, and phages that have departed the biofilm - if they do not re-infect within the next time step - are assumed to be removed by fluid advection. (B) Summary heatmap of the effect of biofilm structure on selection for phage resistance. In the heatmap, simulation outcomes are shown for varying degrees of nutrient availability (which controls the baseline host growth rate) and initial resistant strain frequency. Here both phage mobility and removal rate from the liquid phase are intermediate, and the bacterial fitness cost of phage resistance is 5% of the maximum growth rate (see Figure S1 for extensive exploration of these factors). Resistant cells increase in frequency when initially uncommon (blue squares in heatmap), but when they are initially common, their relative abundance either stays the same (white squares) or decreases (red squares).

To understand the population dynamics of phages in the presence of biofilms that contain both susceptible and resistance bacterial strains, we constrained our simulations using experimentally measured parameters for bacterial growth, phage replication, and nutrient diffusion (See Table S1), based on *E. coli* and its lytic phage T7 (the same species used in our experiments, see below). We explored the impact of factors that are likely to vary in natural environments where phage-biofilm interactions occur. The first is nutrient availability, which controls overall biofilm expansion rate (37, 38). We also varied the initial population ratio of susceptible to resistant host bacteria. In this way, we could test for the invasibility of phage-resistant and phage-susceptible cells when rare. For example, if resistant cells always increase (/decrease) in frequency regardless of their initial fraction, we can infer that they are being positively (/negatively) selected. On the other hand, if they increase when initially rare but decrease when initially common, then we can infer that resistant and susceptible cells will tend toward coexistence (39). We also tested for the effect of variation in the fitness cost of phage resistance, variation in phage diffusivity, variation in how phages were introduced to the biofilm surface, and whether phages were introduced at earlier or later time points during biofilm growth (see Supplementary Information). Importantly we also explored the impact, if any, of the mechanism of phage resistance.

Phage resistance can manifest in different ways; for example, in the case of *E. coli* and the lytic phage T7, which attaches to LPS to initiate infection, the host can evolve resistance by modification or partial loss of the LPS biosynthesis machinery, or by loss of thioredoxin A, which is co-opted and required by T7 as a phage DNA polymerase processivity factor (40). In the case of LPS mutants, phages cannot bind the cell surface, and thus phage-host encounters leave both phage and host intact. Mutants of *E. coli* that have lost thioredoxin A, however, allow phage entry, but not replication, and thus cause an abortive infection in which the host and the phage are both killed. In our experimental tests below, we use the thioredoxin A mutant as our phage-resistant strain, and for correspondence we use for our simulations in the main text a resistant mutant that both neutralizes phages and is neutralized when a phage attaches. However we repeat all simulations for the scenario in which neither the host cell or phage is neutralized by a contact event and phages are free to continue diffusing (such as for surface mutants), and for the scenario in which the bacterial host is unaffected while the phage is neutralized by a contact event (such as for CRISPR-Cas9 based immunity).

### Biofilms robustly facilitate coexistence of phage-resistant and -susceptible cells

The full results of our parameter sweeps are shown in Figures S1 and S2, and for clarity we show a representative sub-set of these results in Figure 1B, where the fitness cost (i.e. growth rate decrement) of displaying phage resistance is a 5% reduction in maximum growth rate, and phages are moderately impeded from diffusion in biofilms. On the scale of the whole biofilm simulation space, the overriding pattern of our simulations was positive selection for phage resistant cells when they are initially rare, and either neutral or negative selection for resistant cells when they are initially common (SI Figure S1; SI Video S1 and Video S2). The only exceptions occur when phage mobility is extremely high, in which case the system behaves as though it were a well-mixed culture and phage resistance is uniformly positively selected (Figures S1); or when phage mobility is so severely constrained that viral particles never have an opportunity to ‘find’ susceptible hosts by diffusion, in which case the phage-resistant and phage-susceptible cells compete solely according to their growth rates (Figures S1). We observed the same qualitative results when our simulations were implemented in 3-D space (SI Video S3 and Video S4). The results were also the same regardless of the mode of resistance among the bacteria; the same trends were upheld if the resistant strain caused abortive infections, if neither resistant hosts nor phages were neutralized by mutual contact, or if phages alone were neutralized by contact with resistant hosts (Figure S2).

The results broadly and robustly support the prediction of coexistence of phage-resistant and phage-susceptible cells in biofilm environments. We observed the same qualitative pattern as shown in Figure 1B when we varied the biofilm size at which phages were introduced, and there was similarly little effect if phages were introduced at a single point or evenly along the entire biofilm surface (Figure S3). The strength of negative frequency-dependent selection, and the predicted stable frequencies of resistance and susceptible cells, are tuned by phage mobility and the cost of phage resistance (15), but the overall qualitative pattern of predicted coexistence is highly robust to parameter changes (Figure S1) (39, 41–43). We next looked for the details of this negative frequency-dependence: why do phage-resistant cells fare well when rare, but fare poorly when common?

#### Clearance of susceptible cells when they are common

When phage-susceptible cells start in the majority within a biofilm, the few resistant cells initially in the population are concentrated into small isolated groups. As a result, when phages enter the system, they have ready access to susceptible hosts that occupy the majority of space, and the propagating infection eliminates most or all of the susceptible population. After this clearance event, the few remaining phage-resistant cells have an abundance of open space to occupy as they continue to grow with reduced competition for nutrient sources in the surrounding medium (Figure 2A,B). Unless the cost of phage resistance is very high (Figure S1), resistant cells tend not to reach fixation due to small pockets of susceptible cells that are protected from phage exposure by neighboring resistant cells (Figure 2B). This latter effect is strengthened if resistant cells are initially abundant, as detailed below.

**Figure 2.**
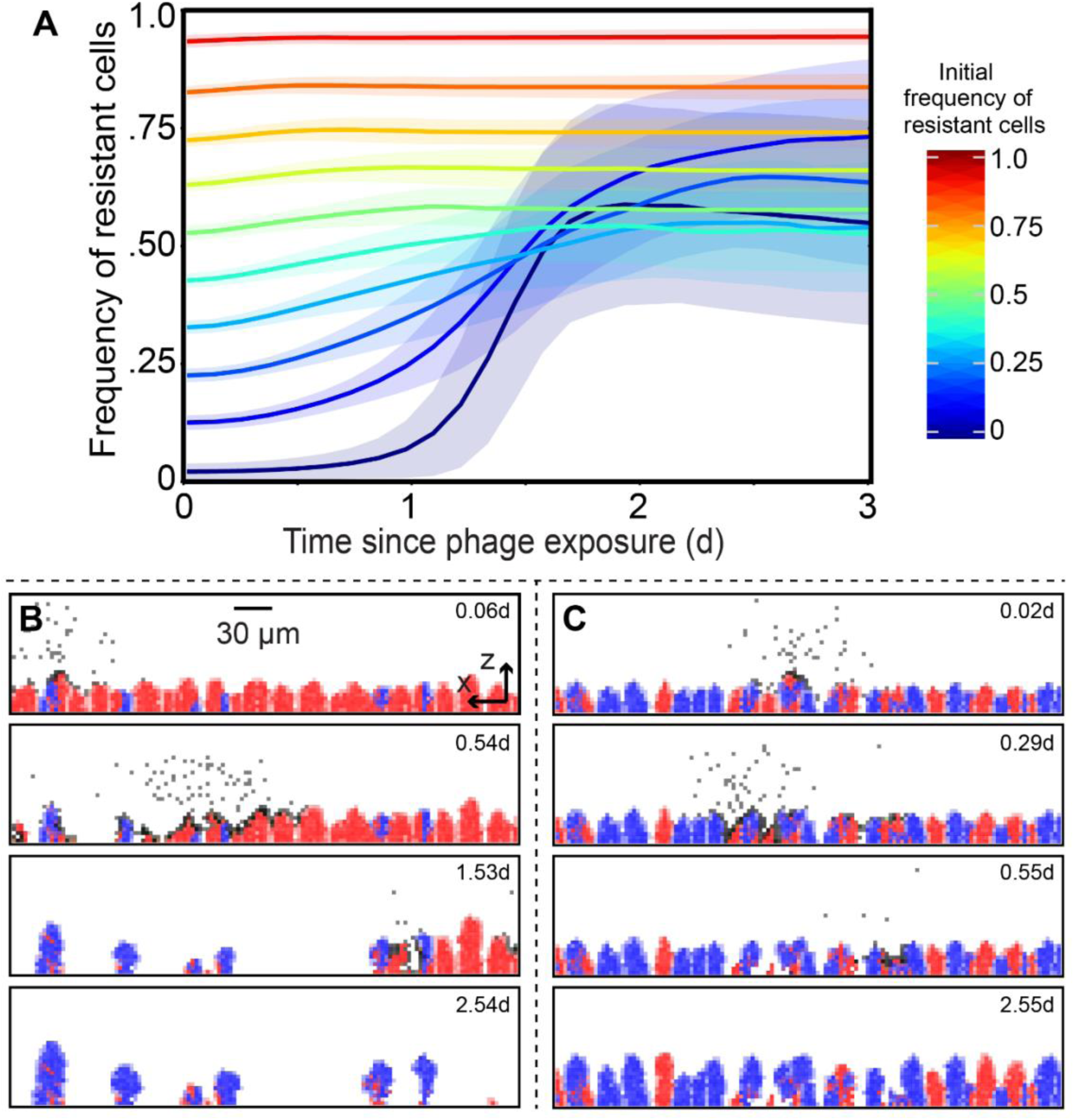
Simulated population dynamics of phage-resistant and susceptible bacteria within biofilms. These dynamics underlie the competition outcomes in Figure 1. (A) The frequency of resistant cells is shown in traces colored according to their initial frequency, with the standard deviation across all replicate runs as transparent blue regions around each trace (*n* = 90-100 replicate simulations per trace). (B) When resistant cells are initially a minority, susceptible cells are exposed to phages and largely killed off, allowing resistant cells to re-seed the population and markedly increase in relative abundance relative to the strain ratio prior to phage exposure. (C) When resistant cells are initially more common, and phages cannot diffuse freely through the biofilm, susceptible cells are spatially protected from phage exposure because phages are sequestered in clusters of resistant cells.

#### Phage sequestering by resistant cells when they are common

When phage-resistant cells are initially common, phage-susceptible cell clusters are isolated among larger groups of resistant cells. If phage diffusion is even moderately impeded by the presence of biofilm, then susceptible cells gain protection from phages. This occurs because phages become trapped on the periphery of clusters of resistant cells, and because phages released into the liquid phase are often blocked from long-range movement by groups of resistant cells in their path. The lower the frequency of susceptible cells in the initial inoculum, the stronger the effect of these spatial phage protection mechanisms. In this scenario, if there is no cost to resistance, then susceptible and resistant cells compete neutrally. If there is a fitness cost to resistance, then susceptible cells have an intrinsic growth rate advantage, and they increase in frequency if they are initially rare (Figure 2A,C).

### Experimental model of phage resistance population dynamics

Our simulation results strongly support the prediction for coexistence of phage-susceptible and phage-resistant cells in biofilm environments, nearly regardless of variation in any major features of the system. Here we set out to test this prediction using an experimental model of biofilm growth under lytic phage attack. Biofilms of *E. coli* were cultivated in microfluidic devices, including co-cultures of wild type AR3110 (WT), which is T7-susceptible, and an isogenic strain harboring a clean deletion of *trxA*, which does not support phage replication (see Materials and Methods). The Δ*trxA* mutant lacks thioredoxin A, which is an essential DNA processivity factor for the lytic phage T7. This deletion mutant therefore causes an abortive infection in which phage attachment occurs and the host is killed, but the phage is not able to replicate or lyse the host (40). We chose the Δ*trxA* mutant as representative of phage-resistant variants because it does not support phage propagation but is able to form biofilms normally. Almost all other mutations conferring T7 resistance are in the LPS assembly machinery, and our pilot experiments indicated that these mutant classes are severely defective for biofilm formation, and so do not allow the experiments described below to be performed. This biofilm defect is a notable fitness cost of LPS-modification-dependent phage resistance, but in order to test our predictions we required a T7-resistant mutant capable of biofilm formation and thus focus on the Δ*trxA* background for the remainder of the paper. Growth curves in shaken liquid media identical to that used for biofilm experiments indicated that the phage-resistant Δ*trxA* mutant has a growth rate cost of 7.9% +/-0.69% (Figure S4)

The *E. coli* experimental biofilms were cultivated in microfluidic devices composed of a chamber molded into PDMS, which was then bonded to a glass coverslip for imaging on an inverted confocal microscope. Prior work has shown that even biofilms of phage-susceptible WT *E. coli* AR3110 can protect themselves from phages after ∼60 h of growth, when they begin to produce a curli amyloid fiber mesh that blocks phage diffusion (23). Here biofilms of WT and Δ*trxA* mutant were cultivated for only 48 hours prior to phage exposure, such that no curli-mediated phage protection could occur during the initial phage exposure. In different runs of the experiment, mimicking our simulation approach, we inoculated the glass bottom of flow devices with varying ratios of phage-susceptible and phage-resistant bacterial cells. Analogous to the simulations, we allowed biofilms to grow undisturbed for 48 hours and then subjected them to a pulse of high-density phage suspensions (Figure S5; Materials and Methods). Biofilm populations were then imaged by confocal microscopy at regular intervals for 2 days. For each imaging session, the entire biofilm volume was captured in successive optical sections.

We found that when phage-resistant cells were initially rare, susceptible cells were killed off by phage exposure and mostly cleared out of the chambers, opening new space into which resistant cells could grow for the remainder of the experiment (Figure 3A,B). As in our simulations, resistant cells often did not reach fixation, as small clusters of susceptible cells remained. On the other hand, when phage-resistant cells were initially common (∼60% of the population, or more), the relative fraction of resistant and susceptible host bacteria did not substantially change following phage treatment (Figure 3A,C). We did not observe localized cycling of resistant and susceptible cells, as one might predict in closed and shaken liquid culture conditions, most likely because phages were either sequestered locally within clusters of resistant cells (Figure 4), or advected out of the system by ongoing fluid flow in our microfluidic devices.

**Figure 3.**
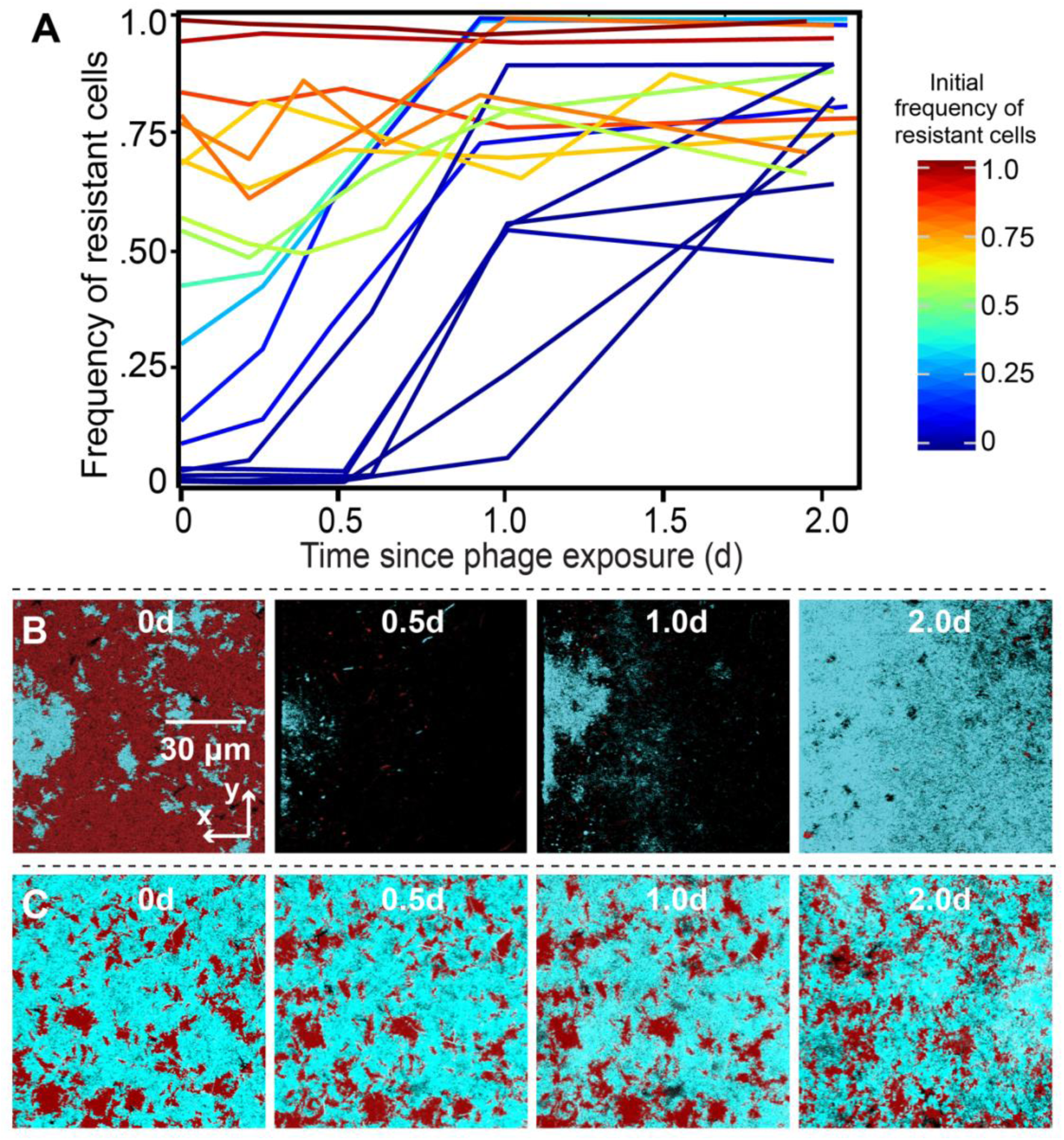
Experimental test of model predictions for phage-biofilm coexistence. Biofilms containing mixtures of phage T7-susceptible AR3110 *E. coli* and a phage T7-resistant mutant carrying a deletion of *trxA* were grown for 48 hours before administering a pulse of phages to the two-strain biofilm population. The frequency of resistant cells is shown in traces colored according to their initial frequency, where each trace is an independent run of the experiment. (A) Population dynamics traces showing the frequency of phage-resistant *E. coli* as a function of its initial population frequency. Each trace is a single replicate of the experiment, with varying initial ratios of the two strains as in our simulations (B, C) Time series of phage-resistant (blue) and phage-susceptible cells (red) following a pulse of phages into the chambers. Panels from left to right show biofilms at ∼ 0, 0.5, 1, and 2 days after phage exposure. Each image is an x-y optical section from a stack of images covering the whole biofilm volume, taken by confocal microscopy.

**Figure 4.**
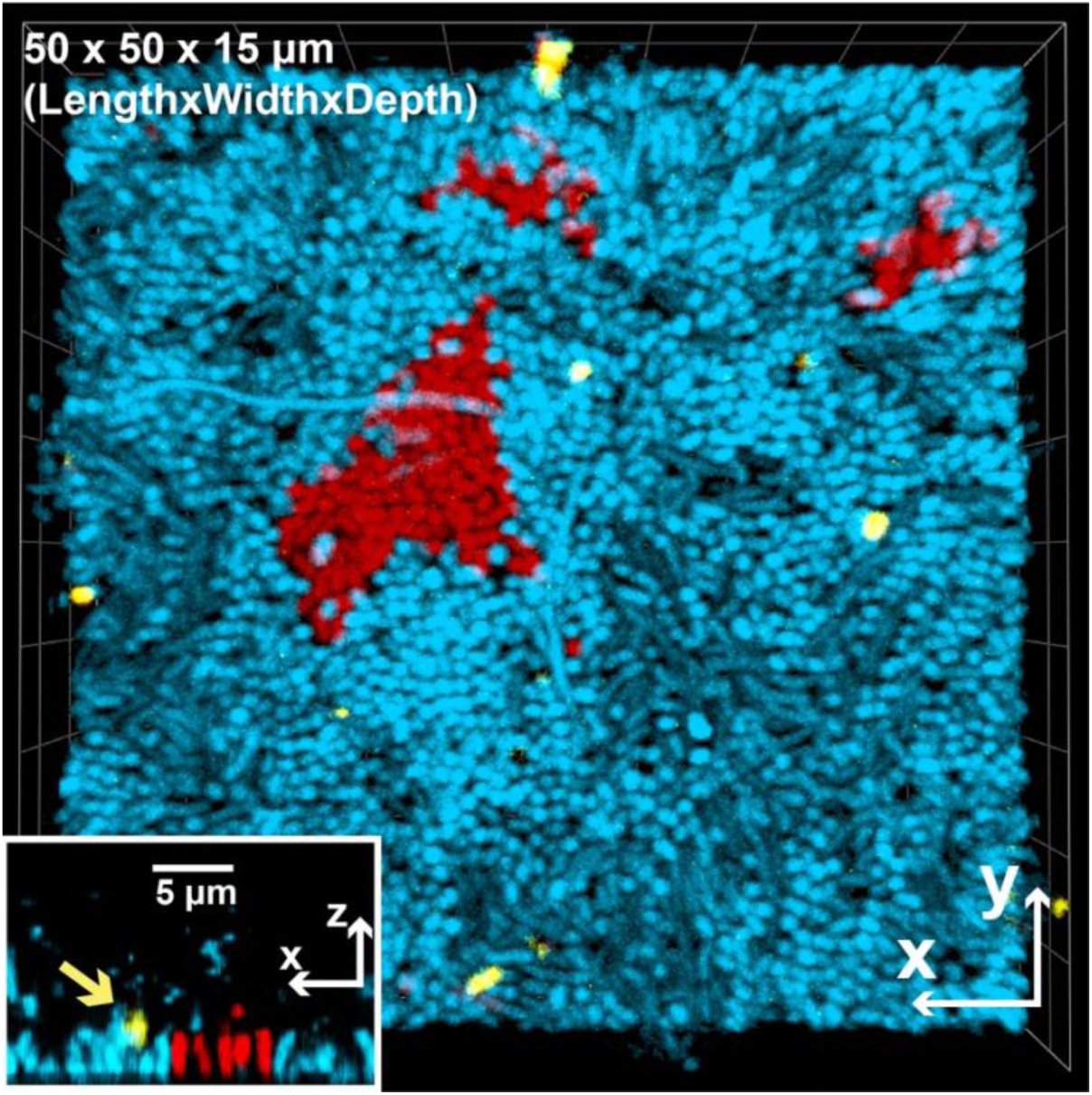
Experimental demonstration of phage sequestration within clusters of phage-resistant bacteria (blue) in a mixed-strain biofilm with phage-susceptible bacteria (red). Purified phages stained with Alexafluor-633 (shown in yellow) were added to 48 h biofilms in which resistant cells were inoculated as 95% of the founding population. The central image is a top-down view of a 3-D rendering measuring 50μm × 50μm × 15 μm [L × W × D]. The inset image is a 2-D projection of a vertical slice through a 3-D volume. The yellow arrow points to an immobilized phage on a cluster of resistant cells. Note that phages are much smaller than the minimum resolvable volume of a confocal fluorescence microscope like that used here; as a result of this effect and the fact that their Alexafluor-633 tag is very bright, the phages appear larger than their true size.

Our experimental results thus displayed a good qualitative match to our models. The spatial patterns underlying these outcomes were the same as those observed in our simulations, including a clearance of susceptible cells when resistant cells are initially rare. In this condition, susceptible cells are exposed to phages; the remaining resistant cell clusters then have ample room to multiply (Figure 3B). Our experiments also confirmed that susceptible cells are protected when they are initially rare: when resistant cells are common, they often sequester phages away from susceptible cells, which then remain near their initial frequency in the population (Figure 3C). To further test this inference, we introduced fluorescently labelled T7 phages to biofilms initiated with a majority of resistant bacteria, and directly observed that these phages immobilized in regions of the biofilm occupied purely by resistant cells (Figure 4, additional replicas in Figure S6).

## Discussion

Our results provide a foundation for understanding how decreased phage mobility and sequestration to resistant hosts within biofilms determine the population dynamics of phage resistance. Using simulations with extensive parameter sweeps, we found a dominating trend toward negative frequency-dependent selection for phage resistance that is remarkably robust to parameter changes. This outcome is matched by our microfluidic model of biofilm formation visualized at single cell resolution, and it reinforces and generalizes predictions from more abstracted models in the literature (14, 16, 29).

The origins of frequency-dependent selection are tied to the cell movement constraints and competition for space in biofilms. When phage-resistant bacteria are initially rare, introduced phages have open access to susceptible hosts, which are mostly killed, leaving empty space for the residual resistant cell clusters to occupy. On the other hand, when phage-resistant bacteria are initially common, they create barriers between phages and clusters of susceptible cells. So long as there is impeded diffusion of phages through the biofilm volume, this spatial arrangement provides protection to susceptible cells, whose population frequency can then drift or increase significantly depending on the fitness costs of phage resistance. On the basis of our parameter sensitivity analyses, we infer that this pattern is an inevitable consequence of the spatial constraints inherent to biofilm communities.

We tested these outcomes experimentally using microfluidic culture and confocal microscopy of mixed *E. coli* biofilms containing resistant and susceptible hosts; these trials gave an excellent qualitative match to the simulations, and we could document both the clearance and phage sequestration effects, depending, as anticipated from simulations, on the initial fractions of resistant and susceptible bacterial cells. Because LPS mutants of *E. coli* appear to be severely impaired for biofilm formation, we were only able to experimentally test the case in which abortive infection is the resistance mechanism. In this manner phages are limited in their diffusion not only because of barriers of resistant cells, but also because of sorptive scavenging; that is, they are sequestered by resistant cells, and both the host and phage are neutralized by the encounter (16, 44–48). We emphasize, however, that our simulations very strongly suggest that even in the event that neither host nor phage is neutralized by a mutual encounter and phages are free to continue diffusing, such as when a mutant gains phage resistance via loss of the cell surface phage receptor, the same pattern of negative frequency-dependent selection for resistance will be upheld in the vast majority of conditions.

Our results also draw an analogy between phage ‘epidemics’ on the sub-millimeter scale of biofilms and the process of herd immunity studied for decades at much larger spatial scales in populations of plants and animals (49–51). When enough of the population is resistant, a spreading pathogen is no longer able to establish sufficient infections to amplify itself, and the susceptible portion of the population is protected (49). These observations in turn have several general implications. We anticipate that the arms race of phage attack and host defense can have a very different landscape in biofilms compared with planktonic populations (2, 5, 7, 14, 52). A rich history of research has shown that phages can rapidly eliminate susceptible host cell populations in mixed liquid culture, leading to strong selection for phage resistance (2–4, 53). In biofilms, by contrast, our results predict widespread and easily maintained polymorphism in phage resistance ability. This kind of standing variation can arise due to minority advantage (i.e., kill-the-winner) mechanisms (54–57), in which phages or other parasites are selected to target the most abundant constituent strains of a population.

The mechanism we describe here is distinct from kill-the-winner based selection, but complementary: susceptible cells in the minority are unlikely to be exposed to phages in the first place, as they are shielded by resistant cells blocking phage diffusion. The arms race between phages and host bacteria, therefore, is likely to take different evolutionary trajectories that move at slower speeds than those typically observed in liquid culture. This outcome echoes results observed in the early phage-host coevolution literature, where it found that for bacteria that form ‘wall populations’ on the inside of shaken liquid culture tubes, phage-susceptible bacteria survive at much higher rates than in the well-mixed planktonic phase (58). These wall populations are now known as biofilms, and here we have directly visualized the spatial protection process that allows susceptible cells to survive where otherwise they would not. The results obtained here also make concrete on the microscopic scale how variable access of phages to susceptible hosts shifts populations to steady states in which phage-resistant and phage-susceptible bacteria should robustly coexist (5, 14).

Our observations also bear on the efficacy of phage therapies, for which one of the most promising potential benefits is selective elimination of target pathogens from a community of otherwise commensal or beneficial microbes (12, 27, 56, 59, 60). This is a particularly compelling advantage relative to broad-spectrum antibiotics that can kill off not just the target pathogen but also many other members of a patient’s microbiota, sometimes with severe side effects (61). Our work suggests that while it might be possible to completely eliminate target bacteria with lytic phages from a mixed population, the success of this approach depends heavily on the community composition and spatial structure. Phage-susceptible cells can be much harder to target and can coexist with resistant cells, or presumably cells of other species that phage cannot target, due to the protective effects of phage sequestration and diffusional blocking. Successful phage treatment will likely depend on disruption of the biofilm architecture to ensure exposure of target bacteria to the therapeutic. It should be noted, however, that our work here only examines two strains of the same species, and whether these conclusions apply to multi-species consortia (62), whose biofilm architectures can differ substantially, is an important topic for further work.

The models developed here do not address the possibility of refuges created by quiescent bacteria in the basal layers of biofilms where nutrients have been depleted (14). This did not appear to be an important feature of our experimental biofilms, which agreed well with simulation predictions. However, quiescent cells could potentially be significant in other conditions, especially for cell groups that accumulate thicker mats with large, nutrient-starved populations in their interior. We also do not implement ongoing mutations in the different bacterial and phage strains residing in biofilms, using instead strains that are fixed in either the phage-susceptible or -resistant state to examine short term population dynamics. Lastly, and importantly, we omitted from our simulations and experiments the possibility of temperate phage infections, in which the phage genome is inserted to the chromosome of the host organism, emerging to replicate and produce new phages when the host is under duress. Temperate phages present a wide diversity of potential outcomes, especially considering that they can impart new phenotypes to their bacterial hosts. Tackling the challenge, both theoretically and experimentally, of how temperate phages enter, alter, and evolve within multispecies microbial communities is an important area for future work.

## Author Contributions

CDN conceived and supervised the project; CDN and VB designed simulations and experiments. MS developed the simulation framework and performed simulation data collection. MCB performed experiments and image processing of microscopy data. MKS, MCB, BK, KD, VB, and CDN analyzed and interpreted data. MKS, MCB, BK, KD, VB, and CDN wrote the paper.

## Acknowledgements

We are grateful to Will Harcombe, Wolfram Möbius, Ben Wucher, Swetha Kasetty, and Sara Mitri for comments on earlier versions of the manuscript. Input from Jim Bull was invaluable in completing the paper. MCB is supported by a GANN Fellowship from Dartmouth College. KD is supported by the European Research Council (StG-716734), the Deutsche Forschungsgemeinschaft (SFB 987), and the Behrens-Weise-Foundation. VB is supported by NSF ABI 1458347 and a UMass President Science and Technology award. CDN is supported by the National Science Foundation MCB 1817342, a Burke Award from Dartmouth College, a pilot award from the Cystic Fibrosis Foundation (STANTO15RO), NIH grant P30-DK117469, and NIH grant P20-GM113132 to the Dartmouth BioMT COBRE.

## Supplemental Information

### Supplemental Materials and Methods

Details of methodology for development of simulation framework and experimental techqnieus used in the paper

### Supplemental Table S1

List of parameters used for simulations, including references where applicable.

### Supplemental Figures 1-6

Additional data in support of the main text.

### Supplemental Videos

**SI Video 1:** A video illustrating the clearance of almost all susceptible cells (red) by phage (black) infection. This occurs when resistant cells (blue) are initially rare in the population. This video is the extended time series from which frames were taken for Figure 1B of the main text.

**SI Video 2:** A video illustrating the sequestration of phages (black) by majority resistant cell clusters (blue), protecting most of the minority susceptible cell population (red) from phage exposure. This occurs when resistant cells are initially common in the biofilm population. This video is the extended time series from which frames were taken for Figure 1C of the main text.

**SI Video 3:** A biofilm simulation in 3-dimensions illustrating the clearance effect by which susceptible cells (red, infected cells shown in green), when common in the biofilm population relative to resistance cells (blue), are mostly or entirely killed off by a propagating phage infection.

**SI Video 4:** A biofilm simulation in 3-dimensions illustrating the phage sequestration effect by which susceptible cells (red, infected cells shown in green), when initially rare in the biofilm population, are protected from phage exposure by the majority of resistant cell clusters (blue) in their surroundings, which prevent phages from reaching susceptible hosts in which to infect and multiply.

## Materials and Methods

### Phage-biofilm modeling simulation framework

The simulation framework used for this study is an updated and expanded version of a modeling approach developed in Simmons et al. (15). The major changes include a new implementation of bacteria as individual particles rather than a homogeneous biomass, and a new implementation of phage diffusion, detailed below. The simulations are built on a grid-based approach for tracking bacteria, phages, and solute concentrations; spatial structure in the system is thus resolved at the level of grid nodes (which are 3μm × 3μm for the simulations described in this paper). Within a grid node, bacteria and phages are tracked individually but assumed to interact randomly. Using the FiPy partial differential equation solver for Python (63), the same grid system is used to calculate nutrient diffusion from a bulk layer above the biofilm toward the cell group surface, where it is consumed by bacteria (35, 37, 64).

As a result of nutrient consumption on the biofilm’s advancing front (Figure 1A) nutrient gradients are created with high nutrient availability in the outer cell layers and lower nutrient availability with increasing depth into the biofilm interior. Cells near the liquid interface grow maximally, while cells deeper in the biofilm interior grow relatively slowly. Fluid flow is modeled implicitly; following prior literature, we allow the biofilm to erode along its outer front at a rate proportional to the square of the distance from the basal substratum (described in detail in Simmons et al. (15)). Further, any phages that depart from the biofilm into the surrounding liquid are advected out of the simulation space within one iteration cycle, which is approximately 7-8 minutes in simulation time (see below).

The simulation framework was written in an object-oriented style. A simulation object is defined *via* the space of the system, number and properties of implemented grid node containers, biological behaviors of bacteria and phages, one-time events (e.g. phage pulse), and simulation exit conditions. Briefly, the space of the system specifies physical information such as physical size and length scale of the grid node array in which cells, phages, and solutes are implemented. The containers hold the information about each modeled individual present in the system. Behaviors describe a container’s interactions with anything else including other containers, space, or time. Events are one-time-use include the inoculation of the system with bacteria or pulses of phages into the simulation space.

Simulations were initiated by first defining the types of container contents, including both bacterial strains/species of interest (phage-susceptible and phage-resistant), phage-infected bacteria, phages, and the growth substrate as a solute. This process includes specifying values for basic biological and physical parameters in the system (e.g. bacterial growth rate, phage infection rate per host-virus contact, phage lag time, phage burst size, nutrient diffusivity, and others; the full list of parameter values and their measurement origins is provided in Table S1). After containers are established in each simulation instance, the simulation proceeds through inoculation of the two bacterial species on the substratum. Phages were not introduced at the outset of simulations but rather at a set time after bacteria were permitted to grow, as described in the main text. Simulations proceed along the following cycle of steps:

1. diffusion of the nutrient substrate,
2. biomass growth and division,
3. lysis of infected bacteria,
4. erosion of biomass,
5. phage movement,
6. detachment of biomass,
7. phage infection,
8. biofilm relaxation (‘shoving’),
9. detachment of bacteriophage.

### Phage mobility implementation

All processes describing phage-bacteria dynamics are equivalent to those presented in Simmons et al. (15) with one exception pertaining to the methods of computing phage entry and exit from the biofilm bacterial volume. This new approach is described in detail below.

Previously, we analytically solved the diffusion equation to approximate the phage density as a function of location in the biofilm. Here, in order to accommodate for possible biological heterogeneity in bacteriophage dynamics (65, 66), we introduced an algorithm for calculating phage movement by modeling each phage’s individual Brownian motion as a random walk. To account for the effect of the biofilm matrix on phage movement, we introduced a new model parameter (the interaction rate, *I*) controlling the diffusivity of phages through areas of simulation space occupied by bacterial biomass (15). We also introduce a rate of removal (*δ*_*p*_) which accounts for the removal of the phage due to the advection of the system during the phage’s motion through the space off of the biofilm, scaling with the square of the distance away from the biofilm. There is an additional implicit advective removal of bacteriophage at the end of the iteration (step 9 above) where any phages remaining off biofilm are removed from the space via advection.

The improved implementation of phage mobility operates as follows. For each phage: We first calculate the number of potential steps that could be taken in the next time interval as: *n* = *D*_*p*_ *dt* / (2 *dl*^2^), and the time of these steps as *dt*_*p*_ = 2 *dl*^2^/ *D*_*p*_, where *dl* is the grid length scale, *D*_*p*_ is the diffusivity of the phage, and *dt* is the simulation time step. Next for each step in *n*: 1) If the phage is off the biofilm, determine whether the phage is removed from advection with probability 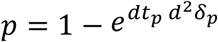, where *d* is the distance away from the biofilm. 2) Next choose a target node by randomly choosing direction. The probability to remain in the current grid node depends on the number of dimensions (See calculation of phage diffusion properties, below)). 3) Determine whether the phage is able to diffuse into the target grid location with probability 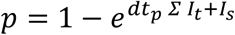, where *I*_*t*_ is the interaction rate at the target grid node, and *I*_*i*_ is the interaction rate at the source node, and we sum over all biomass in those nodes. 4) Finally, if the phage has interacted with biomass, cease motion. If it has not, move the phage to the target grid node. As the interaction rate, *I*, increases, the ability of the phage to diffuse through biomass decreases (e.g., *p* tends to 1), which is a per-individual-phage representation of the phage impedance parameter previously described by Simmons et al. (15). Once the phage stops moving, we evaluate the remaining time as *dt* × *s*/*n*, where *s* is the number of steps taken, from 0 to *n*, and use it in the infection step.

### Calculation of phage diffusion properties

The model for an individual phage taking a step across the grid nodes is that it must diffuse a large enough distance from a grid node. The unnormalized probability density of diffusing within in one place can be described by the solution of the diffusion equation in radial coordinates: 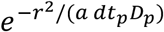. Here *r* indicates the distance away from the starting point, *a* is a constant indicating dimension: *a* = 1 for two dimensions and *a* = 4 for three dimensions, while other terms are explained above. To get the probability of remaining in a radius *ρ*, we integrate from 0 → *ρ* over *r* with a normalization factor which is an integration over all space 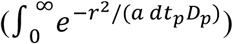. Letting 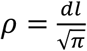 gives a circle whose area is equal to the area of a grid node, and noting that *dt*_*p*_ = 2 *dl*^2^/ *D*_*p*_, the integration yields erf 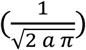, or *p* = 0.42 in two dimension and *p* = 0.22 in three dimensions.

### Details on simulation initial conditions and execution of parameter sweeps

Where possible, biological and physical parameters of the simulation system were constrained according to experimentally measured values for *E. coli* and phage T7, which were the focal species of our experiments as well (see Table S1). Following our previous biofilm dynamics simulation work (15, 38, 67), each simulation starts with an initial ratio of phage-susceptible and -resistant strains on the solid substratum, and these two strains compete for access to space and growth-limiting nutrients that diffuse from a bulk layer above the biofilm. When the biofilm height reaches 30*μm*, (approximately 7 days for the lowest condition and 1 day for the highest), a pulse of bacteriophages to the highest point of susceptible biomass, simulating an individual cell bursting, releasing bacteriophages. Repeating our simulation parameter sweeps with earlier (20*μm* biofilm height) or later (50*μm* biofilm height) phage inoculation had no effect on the qualitative outcomes. Two phage inoculation methods were tested. The first approach to phage inoculation was a 120-virion pulse at a single position at highest point of susceptible biomass in the biofilm. The second was a “spray” of phages in the area just above the biofilm outer surface: 300 phages are added to randomly selected grid nodes 9 *um* above the biofilm. Data reported in Figure 1 correspond to simulations obtained using the first method, but we confirmed that the core results are upheld when using the “spray” method of phage inoculation.

Simulations were run until one of two different exit conditions was reached: either susceptible or resistant cells going to fixation, or the simulation ran to a pre-specified end point (time of infection + 10 days). Simulations were run for 21 different nutrient bulk values corresponding to an approximate time of infection at 1 through 7 days, where the faster growth has slightly greater strain mixing (68, 69). The initial resistant strain frequency also varied from 1% to 99% in 21 steps. Additional simulations were run also for three distinct fitness cost levels of phage resistance, and for three different values of the interaction rate parameter *I*, which effectively varied phage mobility through biofilms easily penetrating the biofilm surface to severely impeded immediately upon biofilm contact (see main text). We ran 100 simulations with different random seeds to completion for each combination of parameters in the main text. Simulations which had a maximum number of phages in any particular iteration of less than 150 were excluded from the analysis, resulting in over 90 simulations for each set in the main text. For the well-mixed control simulations, we disabled all spatially dependent behaviors: substrate diffusion, biomass erosion, biofilm detachment, biofilm relaxation, phage detachment.

### Experimental Materials and Methods

#### Bacterial Strains

Both strains used in this study are *E. coli* AR3110 derivatives, created using the lambda red method for chromosomal modification (70). The Δ*trxA* deletion strain was created by amplifying the locus encoding chloramphenicol acetyltransferase (*cat*) flanked by FRT recombinase sites target sites, using primers with 20bp sequences immediately upstream and downstream of the native *trxA* locus. The FRT recombinase encoded on pCP20 was used to remove the *cat* resistance marker after PCR and sequencing confirmed proper deletion of *trxA*. The wild type *E. coli* AR3110 was engineered to constitutively express the fluorescent protein mKate2, and the *trxA* null mutant was engineered to constitutively produce the fluorescent protein mKO-κ. These fluorescent protein expression constructs were integrated in single copy to the *attB* locus on the chromosome, and they allowed us to visualize the two strains and distinguish them in biofilm co-culture by confocal microscopy.

#### Biofilm growth in microfluidic channels

Microfluidic devices were constructed by bonding poly-dimethylsiloxane (PDMS) castings to size #1.5 36mm × 60mm cover glass (ThermoFisher, Waltham MA) (71, 72). Bacterial strains were grown in 5mL lysogeny broth overnight at 37°C with shaking at 250 r.p.m. Cells were pelleted and washed twice with 0.9% NaCl before normalizing to OD_600_ = 0.2. Strains were combined in varying ratios (see main text) and inoculated into channels of the microfluidic devices. Bacteria were allowed to colonize for 1 hour at room temperature (21-24°C) before providing constant flow (0.1µL/min) of Tryptone broth (10g L-1). Media flow was achieved using syringe pumps (Pico Plus Elite, Harvard Apparatus) and 1mL syringes (25-guage needle) fitted with #30 Cole palmer PTFE tubing (ID = 0.3mm). Tubing was inserted into holes bored in the PDMS with a catheter punch driver.

#### Bacteriophage amplification and purification

T7 phages (23) were used for all experiments. *E coli* AR3110 was used as the phage host for amplification. Purification was conducted according to a protocol developed by Bonilla et al. (73). Briefly, overnight cultures of AR3110 were back diluted 1:10 into 100mL lysogeny broth supplemented with 0.001 M CaCl2 and MgCl2, and incubated for 1 hour at 37°C with shaking; phages from a frozen stock were inoculated and incubated until the culture cleared completely as assessed by eye. Cultures were pelleted, sterile filtered and treated with chloroform. Chloroform was separated from lysate *via* centrifugation and aspiration of supernatant. Phage lysate was then concentrated and cleaned using phosphate buffered saline and repeated spin cycles of an Amicon® Ultra centrifugal filter units with an Ultracel 200kDa membrane (Millipore Sigma, Burlington MA). Purified phages were stored at 4°C.

#### Bacteriophage labeling

Phage labeling began with a high titer phage prep (2×10^10^ PFU/mL) produced using the method described above. 900µL of the phage prep was combined with 90µL sodium bicarbonate (1M, pH = 9.0) and 10µL (1mg/mL) amine reactive Alexa-633 probe (ThermoFisher, Waltham MA) and incubated at room temperature for 1 hour. In this manner the phages were conjugated to dye non-specifically at one or more locations on their capsid coats. Labeled phage were then dialyzed against 1L phosphate buffered saline to remove excess dye using a Float-A-Lyzer®G2 Dialysis Device MWCO 20kD (Spectrum Labs, Rancho Dominguez CA). Labeled phage were diluted in Tryptone broth (10gl-1) to working concentration (2×10^7^ PFU/mL) prior to use.

#### Phage-biofilm microfluidic experiments

Biofilms consisting of varying ratios of susceptible and resistant cells were grown in microfluidic devices for 48 hours at room temperature (21-24°C) under constant media flow (tryptone broth 10gl-1 at 0.1µL/min). Biofilms were imaged immediately prior to phage treatment to establish exact starting ratios of wild type cells (phage-susceptible) and *trxA* deletion mutants (phage-resistant). Subsequently, inlet media tubing was removed from the PDMS microfluidic device and new tubing containing phage diluted in tryptone broth (2×10^7^ PFU/mL at 0.1µL/min) was inserted. Phage treatment continued for 1 hour, after which original tubing was reinserted to resume flow of fresh tryptone broth without phages. Biofilms were imaged approximately 6, 12, 24 and 48 hours after the conclusion of the phage treatment until a population dynamic steady state was reached.

#### Imaging and quantification procedures

Biofilms were imaged using a Zeiss LSM 880 confocal microscope with a C-Apochromat 10X/0.45 water objective or a 40X/1.2 water objective. A 594-nm laser was used to excite mKate2, and a 543-nm laser line was used to excite mKOκ. A 640-nm laser was used to excite Alexafluor 633. Whole chamber Z stacks were acquired by utilizing 1×10 vertical tile scans (total rectangular area ∼500×5000µm). Quantification of biomass was performed using customized scripts in MATLAB (MathWorks Natick, MA) as previously described in Drescher et al. 2014 (74) and Nadell et al. 2015 (75).

**Supplementary Table S1:** Model Parameters used for Simulations.

| Parameter | Value used in the simulations | Description | References where applicable | Representative value ranges and additional references, where applicable |
| --- | --- | --- | --- | --- |
| $x_{max}, y_{max}$ | 900 $\mu m$, 150 $\mu m$ | The physical size of the system | N/A | - |
| $dl, dV$ | 3 $\mu m$, 27 $\mu m^3$ | Length and volume of a grid element | N/A | - |
| $N_{max}$ | 1.1 – 8 $mg L^{-1}$ | Maximum density of substrate (range of values investigated in this study) | (76) | - |
| $N_{max}$ | 0.055 – 0.4 $mg L^{-1}$ | Well-mixed simulation nutrient availability | (77) | - |
| $D_N$ | $2.3 * 10^{-6} cm^2 s^{-1}$ | Diffusivity of substrate | (76) | - |
| $h$ | 15 $\mu m$ | Diffusion boundary layer height | | - |
| $K_N$ | 1.18 $mg L^{-1}$ | Half saturation constant for substrate | (35, 78) | 5 - 225<br>Biofilm heterotrophic bacterial biomass, including fecal coliforms, e.g. *E. coli* (78, 79)<br>4.86<br>*Pseudomonas putida* F1 on glucose (80) |
| $\delta_E$ | 20 $(m h)^{-1}$ | Erosion constant | (36) | - |
| $m_s$ | $10^{-12} g$ | Bacterial mass per cell | (81) | $10^{-12}$<br>*E. coli* DSM 613 |
| $\mu_s (*)$ | 14.1 $day^{-1}$ | Maximum growth rate | (82) | 17.8<br>*E. coli* K-12 on glucose (83)<br>4.8-17.6<br>*E. coli* K-12 on different substrates (84)<br>6.1<br>Wastewater heterotrophic bacterial biomass (85) |
| $S_{max}$ | $200 \text{ g L}^{-1}$ | Maximum active biomass density | (86) | - |
| $Y$ | 0.495 | Yield of substrate converted to biomass | (67) | 0.69-0.77<br>Wastewater bacteria (87)<br>0.41<br>*E. coli* K-12 on glucose (83)<br>0.41-0.51<br>*P. putida* F1 on glucose (80) |
| $\beta$ | 120 | Phage burst size | (8, 88) | Bacteriophage T7 |
| $D_p$ | $3.82 * 10^{-7} \text{ cm}^2 \text{ s}^{-1}$ | Phage diffusivity constant | This Study | Bacteriophage T7 |
| $I$ | $0.067 - 0.12 (m_s \mu\text{m}^3)^{-1} \text{ s}^{-1}$ | Rate of interaction of phage particles with biomass particles | This study | - |
| $\delta_p$ | $0.001 - 10 (\mu\text{m}^2 \text{ h})^{-1}$ | Phage removal rate | (8, 88) | - |
| $\tau$ | 28.8 minutes | Incubation period before lysis | (15) | Bacteriophage T7 |
| $\gamma$ | $2.92 \text{ h}^{-1}$ | Infection rate per biomass per phage | | - |

(\*) The max growth rate is determined from the model equations as $\mu_s = q_s Y$ . $q_s$ is the substrate uptake rate with value $28.5 \text{ g day}^{-1}$ as in Lapidou et al. (82)

## Supplemental Figures

**Figure S1.**
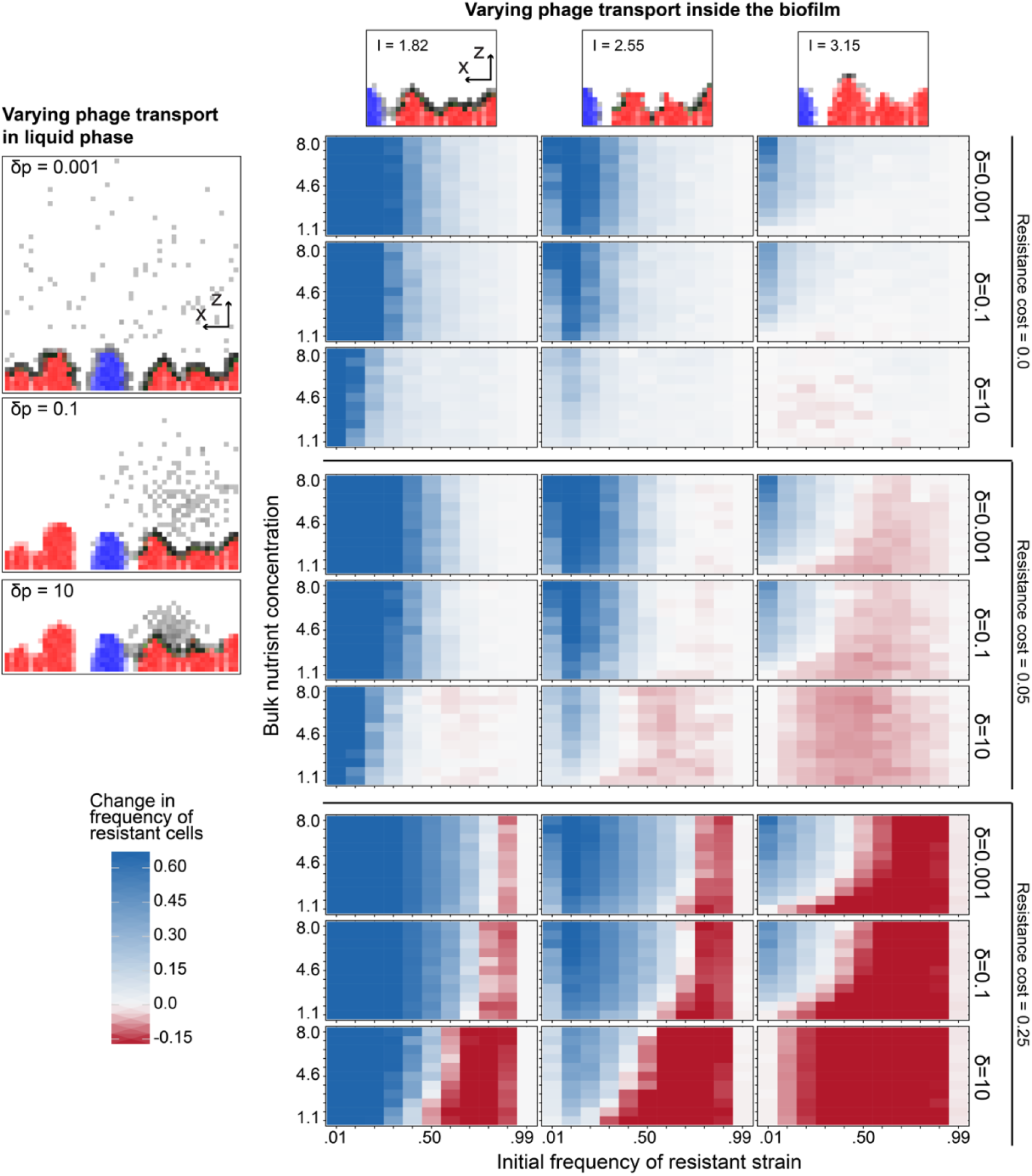
Parameter sensitivity analysis for predicted coexistence of phage-resistant and phage-susceptible cells. The robustness of the predictions outlined in the main text were tested with variation in the cost of phage resistance, the diffusivity of phages through biofilm biomass, and the speed of phage transport/removal in the liquid phase outside of biofilms. As in Figure 1 of the main text, for each parameter combination, simulations were run for a range for varying initial strain frequency, and for varying bulk nutrient concentration, which controls the bacterial growth rate. The heatmaps depict the change in frequency of the resistant strain after phage exposure.

**Figure S2.**
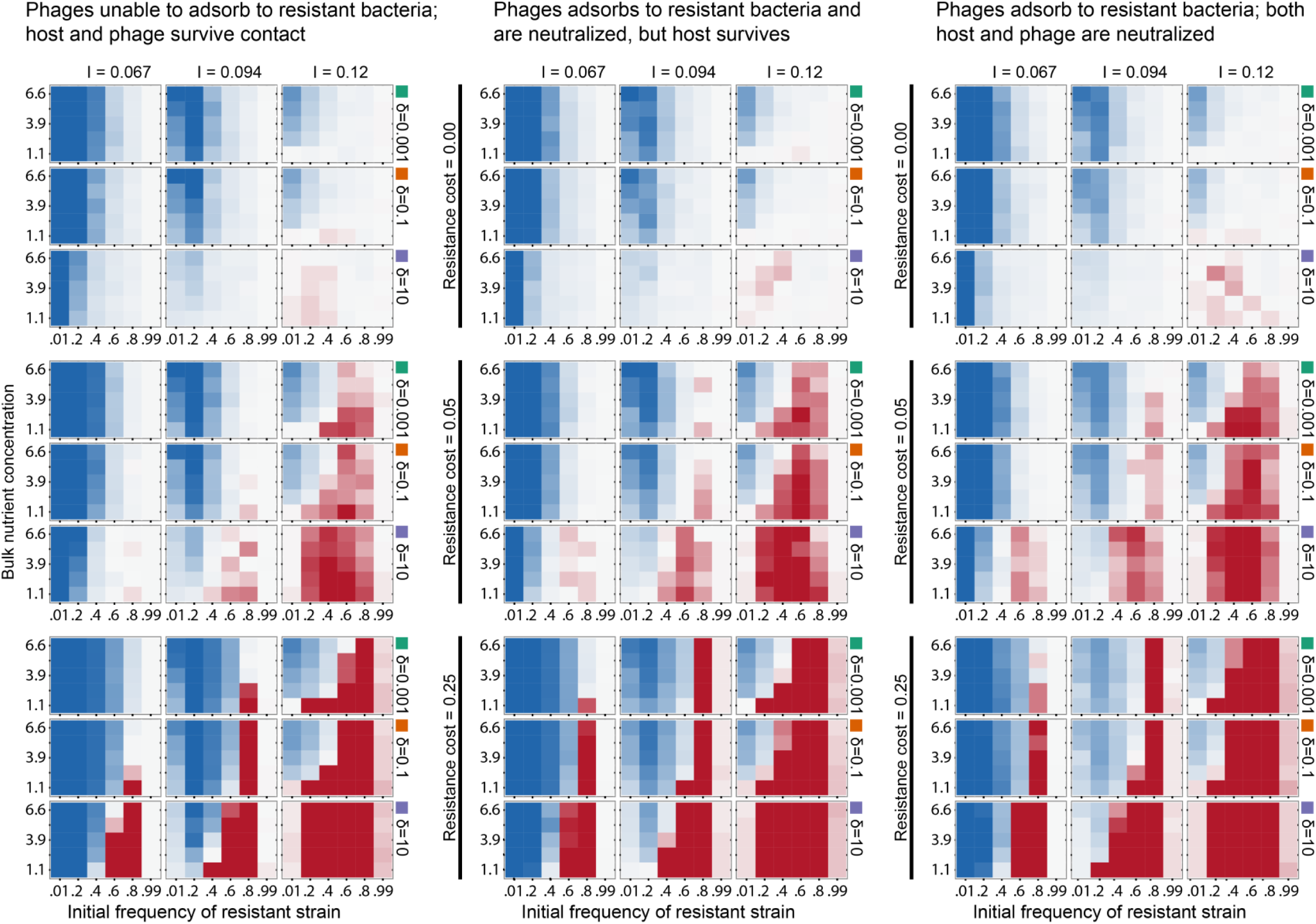
The mechanism of phage resistance does not substantially impact patterns of selection for resistance in biofilms. Here the parameter sweep analysis shown in Figure S1 was repeated (at one quarter resolution) for three different mechanisms of phage resistance on the part of resistant hosts. At left, phages cannot adsorb to hosts, as could be the case for mutants that have lost the phage receptor. In center, phages can adsorb to resistant hosts and re neutralized, but the host survives the encounter and remains viable, as could be the case for CRISPR-Cas9 based resistance to phages. Finally, for comparison we repeat at right the analysis of phage resistance in which both the host and the phage are neutralized by the contact event. This condition represents the abortive infection resistance mechanism that is implemented in our experimental system (see Main Text).

**Figure S3.**
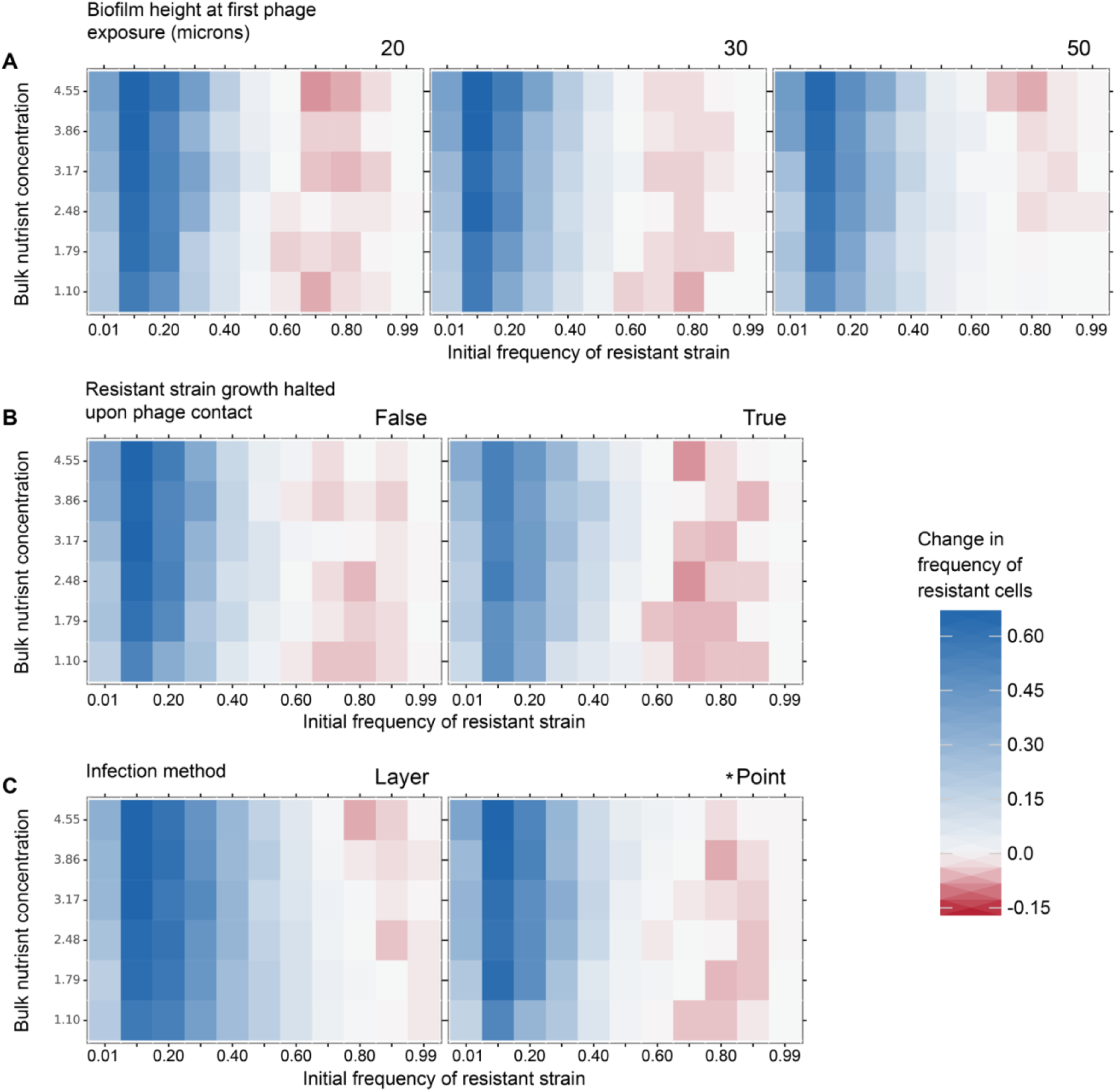
Extended simulations testing the robustness of negative frequency dependent selection for phage resistance. In addition to core simulation parameters assessed in figure S1, we also tested for the robustness of our results against (A) variation in the threshold biofilm height at which phages were pulsed into the system, (B) whether or not resistant cell growth halts upon phage contact, which is the case for some forms of phage resistance that do not permit phage amplification but still allow phage entry into the host cell, and (C) whether phages were introduced in an even layer across the biofilm upper surface, or at a single point on the biofilm surface. All other parameter in these simulations are the same as those used for simulations summarized in Figure 1 of the main text.

**Figure S4.**
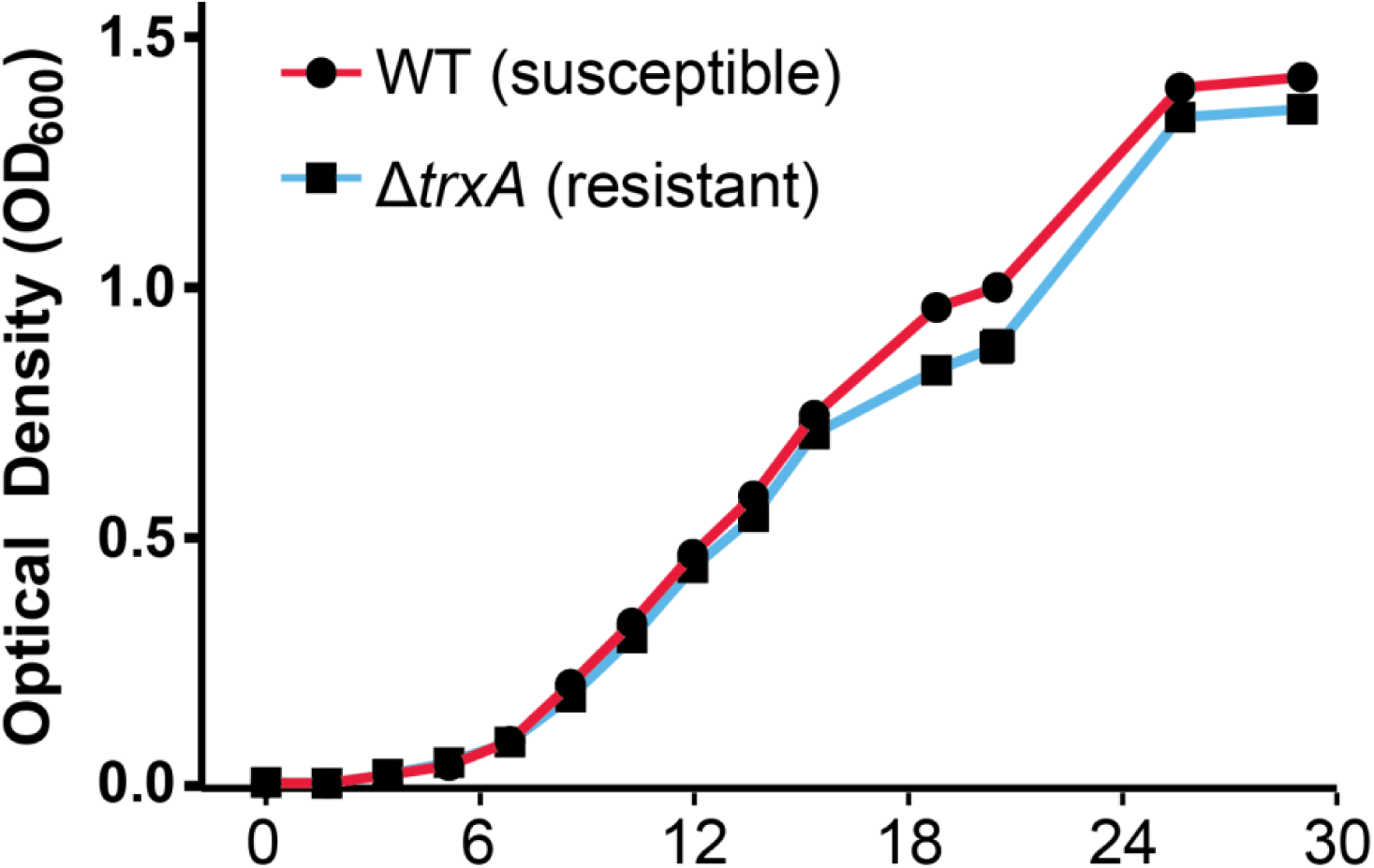
Growth curves of *E. coli* wild type AR3110 (phage T7-susceptible, blue)) and *ΔtrxA* mutants (phage T7-resistant, red) in tryptone liquid culture with shaking at 30°C. Data points denote mean values of 6 total runs of the experiment. Fitting each run to the logistic growth equation yielded an average maximum growth rate of 0.40 +/-0.004 h-1 for the phage-susceptible WT, and a maximum growth rate of 0.37 +/-0.002 h-1 for the phage-resistant *ΔtrxA* mutant.

**Figure S5.**
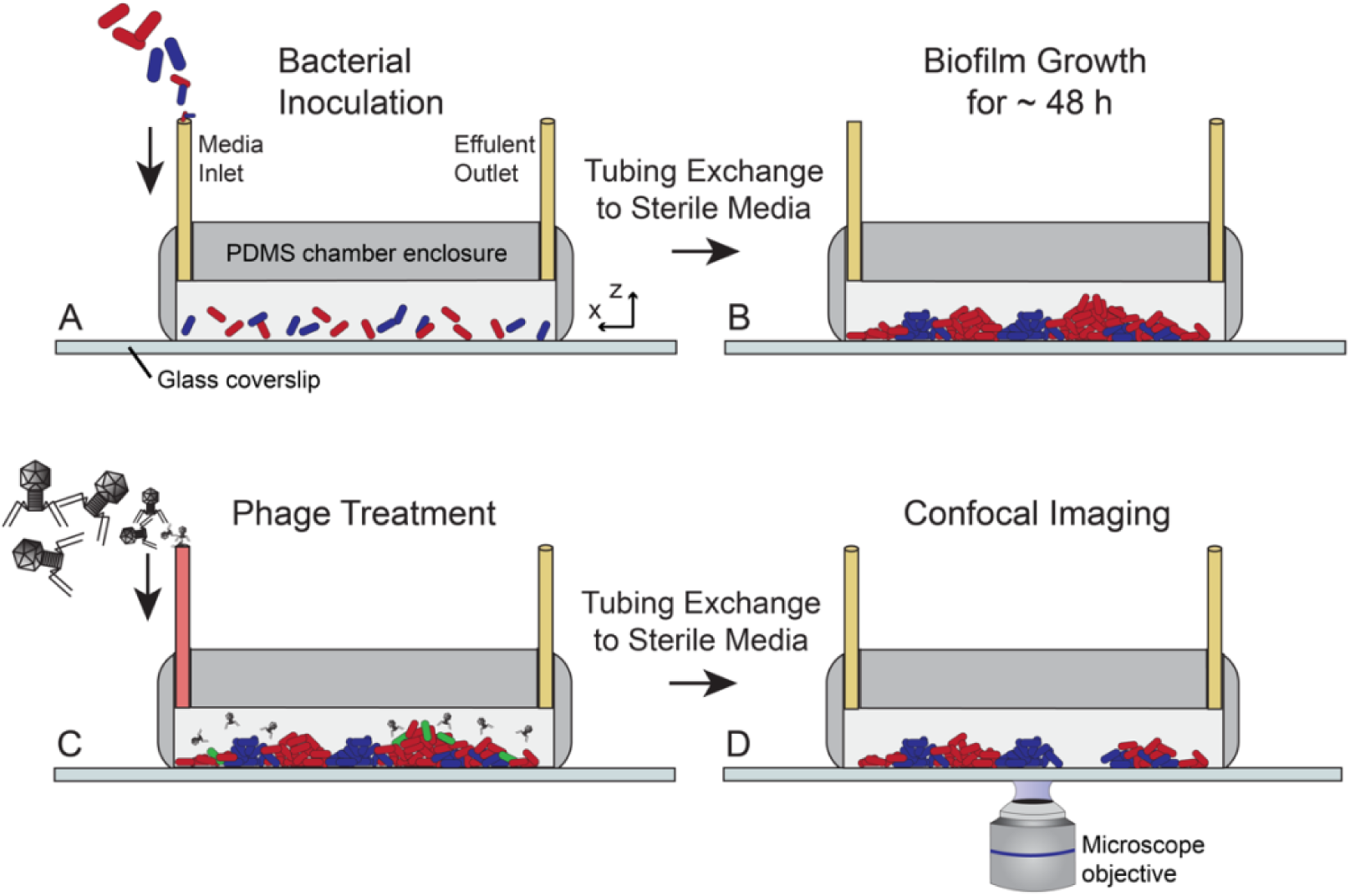
Diagram of experimental biofilm growth and phage treatment regime. (A) Biofilms were grown by inoculating phage-susceptible and -resistant cells in controlled ratios (see main text) onto the glass bottom of PDMS microfluidic devices. B) Biofilms were grown in the absence of phage for 48 hours, after which (C) the medium inlet tubing was switched to perfuse biofilms with T7 phages. (D) the inlet tubing was replaced again to continue flow of fresh media, and image series were acquired by confocal microscopy.

**Figure S6.**
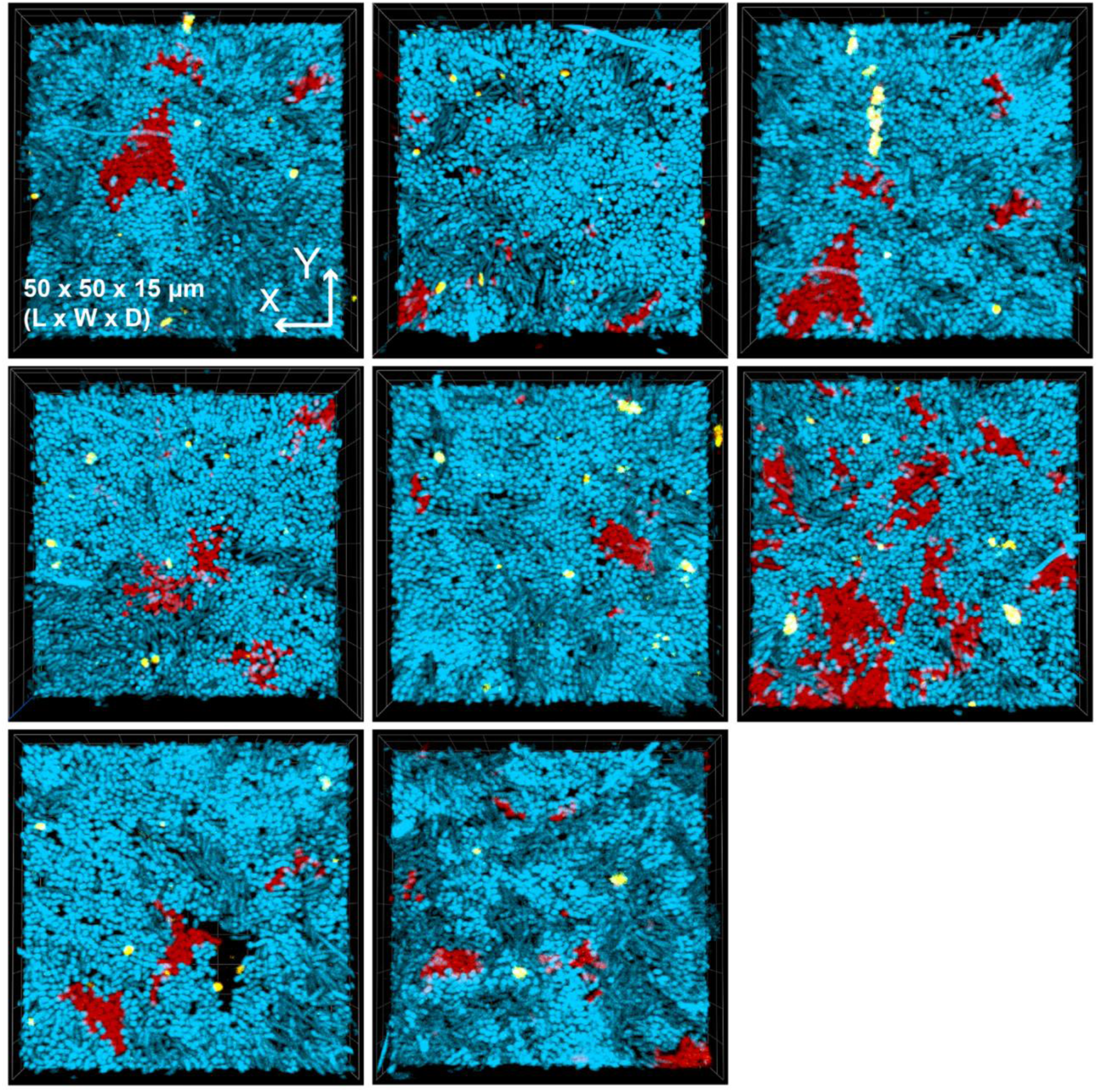
Phages (yellow) trapped by majority resistant bacteria (blue) are unable to reach and infect sparse patches of susceptible cells (red). Additional replicates of the experiment depicted in Figure 4 of the main text, in which biofilms inoculated with 20:1 resistant-susceptible cells were grown for 48 hours and then pulsed with phages for prior to imaging by confocal microscopy. Each panel above is a 3-D biofilm volume rendering ∼ 50μm × 50μm × 15 μm [L × W × D]. Note that the top-left panel is a recapitulation of Figure 4 for comparison with other replicates.

## References

1. Susskind MM, Botstein D (1978) Molecular genetics of bacteriophage P22. Microbiol Rev 42(2):385–413.

2. Koskella B, Brockhurst MA (2014) Bacteria–phage coevolution as a driver of ecological and evolutionary processes in microbial communities. Fems Microbiol Rev 38(5):916–931.

3. Lenski RE, Levin BR (1985) Constraints on the coevolution of bacteria and virulent phage: a model, some experiments, and predictions for natural communities. Am Nat:585–602.

4. Chao L, Levin BR, Stewart FM (1977) A Complex Community in a Simple Habitat: An Experimental Study with Bacteria and Phage. Ecology 58(2):369–378.

5. Brockhurst MA, Buckling A, Rainey PB (2006) Spatial heterogeneity and the stability of host-parasite coexistence. J Evol Biol 19(2):374–379.

6. Harrison E, Laine A-L, Hietala M, Brockhurst MA (2013) Rapidly fluctuating environments constrain coevolutionary arms races by impeding selective sweeps. Proc R Soc B 280(1764):20130937.

7. Brockhurst MA, Buckling A, Rainey PB (2005) The effect of a bacteriophage on diversification of the opportunistic bacterial pathogen, Pseudomonas aeruginosa. Proc R Soc B 272(1570):1385–1391.

8. Abedon ST (2008) Bacteriophage Ecology: Population Growth, Evolution, and Impact of Bacterial Viruses.

9. Abedon ST (2011) Bacteriophages and Biofilms (Nova Science).

10. Labrie SJ, Samson JE, Moineau S (2010) Bacteriophage resistance mechanisms. Nat Rev Micro 8(5):317–327.

11. Samson JE, Magadan AH, Sabri M, Moineau S (2013) Revenge of the phages: defeating bacterial defences. Nat Rev Micro 11(10):675–687.

12. Levin BR, Bull JJ (2004) Population and evolutionary dynamics of phage therapy. Nat Rev Microbiol 2(2):166–173.

13. Weitz JS, et al. (2013) Phage–bacteria infection networks. Trends Microbiol 21(2):82–91.

14. Heilmann S, Sneppen K, Krishna S (2012) Coexistence of phage and bacteria on the boundary of self-organized refuges. Proc Natl Acad Sci U S A 109(31):12828–12833.

15. Simmons M, Drescher K, Nadell CD, Bucci V (2017) Phage mobility is a core determinant of phage–bacteria coexistence in biofilms. Isme J 12:531–543.

16. Bull JJ, et al. (2018) Phage-Bacterial Dynamics with Spatial Structure: Self Organization around Phage Sinks Can Promote Increased Cell Densities. Antibiotics 7(1):8.

17. Eriksen RS, Svenningsen SL, Sneppen K, Mitarai N (2018) A growing microcolony can survive and support persistent propagation of virulent phages. Proc Natl Acad Sci U S A 115(2):337–342.

18. Stewart PS, Franklin MJ (2008) Physiological heterogeneity in biofilms. Nat Rev Microbiol 6(3):199–210.

19. Stewart PS (2012) Mini-review: Convection around biofilms. Biofouling 28(2):187–198.

20. Mah T-FC, O’Toole GA (2001) Mechanisms of biofilm resistance to antimicrobial agents. Trends Microbiol 9(1):34–39.

21. Tseng BS, et al. (2013) The extracellular matrix protects Pseudomonas aeruginosa biofilms by limiting the penetration of tobramycin. Environ Microbiol 15(10):2865–2878.

22. Darch SE, et al. (2017) Phage Inhibit Pathogen Dissemination by Targeting Bacterial Migrants in a Chronic Infection Model. MBio 8(2):e00240–17.

23. Vidakovic L, Singh PK, Hartmann R, Nadell CD, Drescher K (2018) Dynamic biofilm architecture confers individual and collective mechanisms of viral protection. Nat Microbiol 3:26–31.

24. Chaudhry WN, et al. (2019) Mucoidy, a general mechanism for maintaining lytic phage in populations of bacteria. bioRxiv 775056.

25. Scanlan PD, Buckling A (2012) Co-evolution with lytic phage selects for the mucoid phenotype of Pseudomonas fluorescens SBW25. ISME J 6(6):1148–1158.

26. Abedon ST, Thomas-Abedon C (2010) Phage Therapy Pharmacology. Curr Pharm Biotechnol 11(1):28–47.

27. Chan BK, Abedon ST (2012) Phage Therapy Pharmacology: Phage Cocktails. Advances in Applied Microbiology, Vol 78, eds Laskin AI, Sariaslani S, Gadd GM, pp 1–23.

28. Azeredo J, Sutherland IW (2008) The use of phages for the removal of infectious biofilms. Curr Pharm Biotechnol 9(4):261–266.

29. Sutherland IW, Hughes KA, Skillman LC, Tait K (2004) The interaction of phage and biofilms. Fems Microbiol Lett 232(1):1–6.

30. Chaudhry WN, et al. (2018) Leaky resistance and the conditions for the existence of lytic bacteriophage. PLOS Biol 16(8):e2005971.

31. Bull JJ, Vegge CS, Schmerer M, Chaudhry WN, Levin BR (2014) Phenotypic Resistance and the Dynamics of Bacterial Escape from Phage Control. PLoS One 9(4):e94690.

32. Simmons M, Bucci V, Nadell C (2019) SimBiofilm: A framework for indidivual-based biofilm modeling with bacteriophage infection. Available at: https://github.com/simbiofilm/simbiofilm.

33. Nadell CD, Drescher K, Foster KR (2016) Spatial structure, cooperation, and competition in bacterial biofilms. Nat Rev Microbiol 14:589–600.

34. Picioreanu C, Xavier JB, van Loosdrecht MCM (2005) Advances in mathematical modeling of biofilm structure. Biofilms 1(04):337–349.

35. Xavier JB, Picioreanu C, van Loosdrecht MCM (2005) A framework for multidimensional modelling of activity and structure of multispecies biofilms. Environ Microbiol 7(8):1085–1103.

36. Xavier JB, Picioreanu C, van Loosdrecht M (2005) A general description of detachment for multidimensional modelling of biofilms. Biotechnol Bioeng 91(6):651–669.

37. Picioreanu C, van Loosdrecht MCM, Heijnen JJ (1998) A new combined differential-discrete cellular automaton approach for biofilm modeling: Application for growth in gel beads. Biotechnol Bioeng 57(6):718–731.

38. Nadell CD, et al. (2013) Cutting through the complexity of cell collectives. Proc R Soc B 280(1755):20122770.

39. Siepielski AM, McPeek MA (2010) On the evidence for species coexistence: a critique of the coexistence program. Ecology 91(11):3153–64.

40. Qimron U, Marintcheva B, Tabor S, Richardson CC (2006) Genomewide screens for Escherichia coli genes affecting growth of T7 bacteriophage. Proc Natl Acad Sci 103(50):19039–19044.

41. MacArthur R (1972) Geographical Ecology (Princeton University Press, Princeton, NJ).

42. Levin SA (1970) Community Equilibria and Stability, and an Extension of the Competitive Exclusion Principle. Am Nat 104(939):413–423.

43. Chesson P (2000) Mechanisms of Maintenance of Species Diversity. Annu Rev Ecol Syst 31(1):343–366.

44. Yin J, McCaskill J (1992) Replication of viruses in a growing plaque: a reaction-diffusion model. Biophys J 61:1540–1549.

45. Hewson I, Fuhrman J (2003) Vibriobenthos production and virioplankton sorptive scavening. Microb Ecol 46:337–347.

46. Rabinovitch A, Aviram I, Zaritsky A (2003) Bacterial debris - an ecological mechanism for coexistence of bacteria and their viruses. J Theor Biol 224:377–383.

47. Abedon ST (2017) Phage “delay” toward enhancing bacterial escape from biofilms: a more comprehensive way of viewing resistance to bacteriophages. AIMS Microbiol 3(2):186–226.

48. Abedon ST (2016) Bacteriophage exploitation of bacterial biofilms: phage preference for less mature targets? Fems Microbiol Lett 363(3):fnv246.

49. Metcalf CJE, Ferrari M, Graham AL, Grenfell BT (2015) Understanding Herd Immunity. Trends Immunol 36(12):753–755.

50. Levin SA, Durrett R (1996) From individuals to epidemics. Philos Trans R Soc B Biol Sci 351(1347):1615–1621.

51. Levin SA (1992) The Problem of Pattern and Scale in Ecology: The Robert H. MacArthur Award Lecture. Ecology 73(6):1943–1967.

52. Davies E V, et al. (2016) Temperate phages both mediate and drive adaptive evolution in pathogen biofilms. Proc Natl Acad Sci 113(29):8266–8271.

53. Levin BR, Stewart FM, Chao L (1977) Resource-Limited Growth, Competition, and Predation: A Model and Experimental Studies with Bacteria and Bacteriophage. Am Nat 111(977):3–24.

54. Díaz-Muñoz SL, Koskella B (2014) Bacteria-phage interactions in natural environments. Adv Appl Microbiol 89(135):10.1016.

55. Koskella B, Thompson JN, Preston GM, Buckling A (2011) Local biotic environment shapes the spatial scale of bacteriophage adaptation to bacteria. Am Nat 177(4):440–451.

56. Koskella B, Meaden S, Koskella B, Meaden S (2013) Understanding Bacteriophage Specificity in Natural Microbial Communities. Viruses 5(3):806–823.

57. Kunin V, et al. (2008) A bacterial metapopulation adapts locally to phage predation despite global dispersal. Genome Res 18(2):293–7.

58. Schrag SJ, Mittler JE (1996) Host-Parasite Coexistence: The Role of Spatial Refuges in Stabilizing Bacteria-Phage Interactions. Am Nat 148(2):348–377.

59. Levin BR, Bull JJ (1996) Phage therapy revisited: The population biology of a bacterial infection and its treatment with bacteriophage and antibiotics. Am Nat 147(6):881–898.

60. Sillankorva S, Neubauer P, Azeredo J (2010) Phage control of dual species biofilms of Pseudomonas fluorescens and Staphylococcus lentus. Biofouling 26(5):567–575.

61. Buffie CG, et al. (2012) Profound Alterations of Intestinal Microbiota following a Single Dose of Clindamycin Results in Sustained Susceptibility to Clostridium difficile-Induced Colitis. Infect Immun 80(1):62–73.

62. Harcombe WR, Bull JJ (2005) Impact of phages on two-species bacterial communities. Appl Environ Microbiol 71(9):5254–9.

63. Guyer JE, Wheeler D, Warren JA (2009) FiPy: Partial Differential Equations with Python. Comput Sci Eng 11(3):6–15.

64. Lardon LA, et al. (2011) iDynoMiCS: next-generation individual-based modelling of biofilms. Environ Microbiol 13(9):2416–2434.

65. Bucci V, Hoover S, Hellweger FL (2012) Modeling Adaptive Mutation of Enteric Bacteria in Surface Water Using Agent-Based Methods. Water, Air, Soil Pollut 223(5):2035–2049.

66. Hellweger FL, Bucci V (2009) A bunch of tiny individuals-Individual-based modeling for microbes. Ecol Modell 220(1):8–22.

67. Bucci V, Nadell CD, Xavier JB (2011) The evolution of bacteriocin production in bacterial biofilms. Am Nat 178(6):E162–E173.

68. Nadell CD, Foster KR, Xavier JB (2010) Emergence of spatial structure in cell groups and the evolution of cooperation. PLoS Comput Biol 6(3):e1000716.

69. Hallatschek O, Hersen P, Ramanathan S, Nelson DR (2007) Genetic drift at expanding frontiers promotes gene segregation. Proc Natl Acad Sci USA 104(50):19926–19930.

70. Datsenko KA, Wanner BL (2000) One-step inactivation of chromosomal genes in Escherichia coli K-12 using PCR products. Proc Natl Acad Sci U S A 97(12):6640–5.

71. Weibel DB, DiLuzio WR, Whitesides GM (2007) Microfabrication meets microbiology. Nat Rev Microbiol 5(3):209–218.

72. Sia SK, Whitesides GM (2003) Microfluidic devices fabricated in poly(dimethylsiloxane) for biological studies. Electrophoresis 24:3563–3576.

73. Bonilla N, et al. (2016) Phage on tap–a quick and efficient protocol for the preparation of bacteriophage laboratory stocks. PeerJ 4:e2261.

74. Drescher K, Nadell CD, Stone HA, Wingreen NS, Bassler BL (2014) Solutions to the Public Goods Dilemma in Bacterial Biofilms. Curr Biol 24(1):50–55.

75. Nadell CD, Drescher K, Wingreen NS, Bassler BL (2015) Extracellular matrix structure governs invasion resistance in bacterial biofilms. ISME J 9:1700–1709.

76. McCarty PL (2012) Environmental biotechnology: principles and applications (Tata McGraw-Hill Education).

77. Stewart P (2003) Diffusion in Biofilms. J Bacteriol 185(5):1485–1491.

78. Henze M, Grady Jr CPL, Gujer W, Marais GVR, Matsuo T (1987) Activated sludge model no. 1: Iawprc scientific and technical report no. 1. IAWPRC, London.

79. Henze M, et al. (1999) Activated Sludge Model No.2d, ASM2d. Water Sci Technol 39(1):165–182.

80. Oliveira CS, et al. (2009) Determination of kinetic and stoichiometric parameters of Pseudomonas putida F1 by chemostat and in situ pulse respirometry. Chem Prod Process Model 4(2).

81. Loferer-Krößbacher M, Klima J, Psenner R (1998) Determination of Bacterial Cell Dry Mass by Transmission Electron Microscopy and Densitometric Image Analysis. Appl Environ Microbiol 64(2):688–694.

82. Laspidou CS, Rittmann BE (2002) Non-steady state modeling of extracellular polymeric substances, soluble microbial products, and active and inert biomass. Water Res 36(8):1983–1992.

83. Narang A, Konopka A, Ramkrishna D (1997) New patterns of mixed-substrate utilization during batch growth of Escherichia coli K12. Biotechnol Bioeng 55(5):747–757.

84. Beg QK, et al. (2007) Intracellular crowding defines the mode and sequence of substrate uptake by Escherichia coli and constrains its metabolic activity. Proc Natl Acad Sci 104(31):12663–12668.

85. Trojanowicz K, Styka W, Baczynski T (2009) Experimental determination of kinetic parameters for heterotrophic microorganisms in biofilm under petrochemical wastewater conditions. Polish J Environ Stud 18(5).

86. Laspidou CS, Rittmann BE (2004) Modeling the development of biofilm density including active bacteria, inert biomass, and extracellular polymeric substances. Water Res 38(14–15):3349–3361.

87. Esquivel-Rios I, et al. (2014) A microrespirometric method for the determination of stoichiometric and kinetic parameters of heterotrophic and autotrophic cultures. Biochem Eng J 83:70–78.

88. Abedon ST (2009) Kinetics of Phage-Mediated Biocontrol of Bacteria. Foodborne Pathog Dis 6(7):807–815.

